# Expanded diversity of *tfdA* harboring bacteria across the natural and built environment

**DOI:** 10.1101/2022.09.28.509959

**Authors:** Amber M. White, Amarilys Gonzalez Vazquez, Elizabeth A. McDaniel, Benjamin D. Peterson, Paul Koch, Christina K. Remucal, Katherine D. McMahon

**Author notes:** Asterisk indicates co-first authorship of AMW and AGV. Corresponding author address: Amber M. White: 1550 Linden Drive, Madison WI 53706; Twitter: @ambermwhite16.

## Abstract

2,4-Dichlorophenoxyacetic acid (2,4-D) is an herbicide commonly used in aquatic and terrestrial environments that is degraded by bacteria through the TFD pathway. Previous work has relied on culture-based methods to develop primers for qPCR analysis of the gene cassette in environmental samples. In this study, we combined molecular and genomic approaches to examine the accuracy of established *tfdA* qPCR primers on environmental samples and update the phylogeny of *tfdA* genes detected in bacterial genomes. We found most putative 2,4-D degraders are within the Proteobacteria but also found several novel degraders including members of the phyla *Candidatus* Rokubacteria and *Candidatus* Eremiobacteraeota. *In silico* analysis of established primers showed potential amplification of < 5% of putative degrader sequences but 52-100% of experimentally verified degraders when allowing for three and one mismatches between template and primer sequences, respectively. Overall, our work expands the diversity of putative 2,4-D degraders and demonstrates the limitations of culture-based tools for investigating functional diversity of microorganisms in the environment.

**Importance:** Cultivation-based methods can misrepresent the diversity of environmental microorganisms. Our work showcases one example of how culture-based development of molecular tools underestimates the full spectrum of 2,4-D degrading microorganisms. Accurately identifying microorganisms with 2,4-D degradation potential is crucial for understanding the biodegradation potential of a commonly used herbicide across terrestrial, aquatic, and subsurface environments. Additionally, this work reinforces well-documented pitfalls associated with relying on cultured representatives when constructing primers and the challenges of translating findings from a few cultured representatives to understudied or unknown microorganisms in complex environments.

## Introduction

The herbicide 2,4-dichlorophenoxyacetic acid (2,4-D) is used extensively to control invasive and nuisance plants in terrestrial and aquatic environments (1–4) but is a chemical of concern due to its toxicity and potential non-target effects (5, 6). 2,4-D is degraded by bacteria harboring the *tfd* gene cassette, named after the 2,4-D compound (TFD) (7–11). Biodegradation is initiated by *tfdA* gene product, which encodes an alpha-ketoglutarate dependent oxygenase that uses Fe(II) to catalyze the reaction into 2,4-dichlorophenol, carbon dioxide, and succinate (12). Complete mineralization of 2,4-D proceeds via the associated *tfdBCDEF* gene products, which catalyze the transformation of 2,4-dichlorophenol into 3,5-dichlorocatechol; this product is eventually shuttled into the citric acid cycle (13). The reaction catalyzed by the *tfdA* gene product is considered the rate-limiting step of biodegradation because it is transcribed separately from *tfdBCDEF*. There are also generally two copies of the *tfdBCDEF* gene cluster but only one copy of the *tfdA* gene (14). This duplication is likely due to the need for rapid degradation of 2,4-D transformation products that are toxic to bacteria (13). As one of the most used and globally distributed herbicides (11, 15–19), understanding the diversity and prevalence of bacteria with the potential to degrade 2,4-D is critical for understanding the fate and persistence of 2,4-D in the environment.

The *tfd* gene is most often detected in plasmids (9, 13, 20, 21) but can occasionally be found on the chromosome (12–14, 22, 23). *tfdA*-dependent 2,4-D degradation via the TfdA enzyme has been extensively studied on plasmids in *Cupriavidus pinatubonesis*, previously named *Cupriavidus necato*r, *Alcaligenes eutrophus*, and *Ralstonia eutropha* (12, 20, 24, 25). The gene is 819 bp long and is well documented with strains belonging to Alpha, Beta, and Gammaproteobacteria isolated from soil systems (21, 25–27), including several strains capable of using 2,4-D as their sole carbon source (9, 10, 27). There are three recognized *tfdA* gene classes, but these classes are distributed throughout different bacterial phyla because the gene can be horizontally transferred (28). Class I is located on transmissible plasmids and has been detected in Betaproteobacteria. Class II has 76% nucleotide identity to class I, has also been mostly detected in Betaprotebacteria (*Burkholderia* strains), and can also be found on the chromosome where it is referred to as *tfdAα* (22, 23). Lastly, class III has 77% and 80% nucleotide identity to class I and class II, respectively, and has been detected in Beta- and Gammaproteobacteria (28). Phylogenetic analysis of *tfdA* genes sequenced from 31 confirmed 2,4-D degraders found that while there is diversity among the *tfdA* sequences that can be attributed to the different gene classes, gene sequences within the classes are relatively well conserved (25).

Previously, a set of qPCR primers were developed for the quantitative detection of *tfdA* in pure cultures and from the environment (29). Two primer sets that amplify an 81 or 215 bp fragment are expected to amplify all three gene classes. These primers have been applied with variable success in soil (20, 30), sediments (31), aquifer solids (32), and glacial ice (33), but have been infrequently applied in aquatic environments (4, 34, 35). Despite a wide range of environmental applications, the qPCR primers were developed from cultured representatives of soil environments that may miss other environmental representatives. Additionally, no previous studies have applied metagenomic analyses to the *tfdA* gene to the best of our knowledge, suggesting the known diversity of the gene at present is limited to what has been detected through cultivation-based approaches, potentially misrepresenting the complete diversity 2,4-D degraders.

In this study, we combined molecular and metagenomic approaches to examine the accuracy of established *tfdA* qPCR primers on environmental samples and update the *tfdA* phylogeny. Established *tfdA* primers failed to amplify clean PCR products when applied to environmental samples known to degrade 2,4-D. We also identified over 1000 genomes containing the *tfdA* marker from publicly available isolate genomes and metagenome-assembled genomes (MAGs) belonging to over a dozen different phyla from several different environmental sources. Importantly, we found several putative 2,4-D degraders belonging to phyla that, to our knowledge, have not previously been characterized as 2,4-D degraders. Lastly, an *in silico* PCR analysis predicted extremely poor amplification of *tfdA* gene fragments from environmental samples, while genomes of cultured and experimentally verified representatives were indeed predicted to yield PCR products. This work underscores the pitfalls associated with relying on cultured representatives when constructing primers and the challenges of translating findings from a few cultured representatives to understudied or unknown microorganisms in complex environments.

## Results and Discussion

### Confirming specificity of 81 and 215 bp primers in *tfdA* reference genes

Amplification of the standard template for all three gene classes using the 215 bp primer produced products of expected size when visualized on a 1.5% agarose gel (**Figure S1**). Analysis with BLASTN versus the NCBI database also confirmed the sequence of the *tfdA* gene product using the 215 bp primers (**Supplementary Data**). However, initial efforts to amplify *tfdA* using both 81 and 215 bp primers in sediment and soil samples previously exposed to and known to degrade 2,4-D (**Table 1**) yielded non-specific products from extracted sample DNA as evidenced by smearing on agarose gels, even though the positive control produced products of expected size (**Figure 1, Figure 2**). We first hypothesized that co-extracted constituents were interfering with PCR and investigated further amplification interference with two experiments.

**Table 1.** Characteristics of four environmental samples used in TOPO cloning including environmental origin, number of colonies sequenced from each sample, and primer used for PCR amplification. Soil samples described in (77) and water/sediment samples described in (4).

| Environment | Experimental 2,4-D half-life (days) | Colonies sampled | Primer used for PCR |
| --- | --- | --- | --- |
| Soil | < 7 | S | 81 bp |
| Soil | < 7 | T,U,V,W | 81 bp |
| Water | 24 | A,D,E,F,G,H,I,J,K,L,M,N,O | 215 bp |
| Sediment | 24 | B,C,P,Q,R | 215 bp |

**Figure 1.**
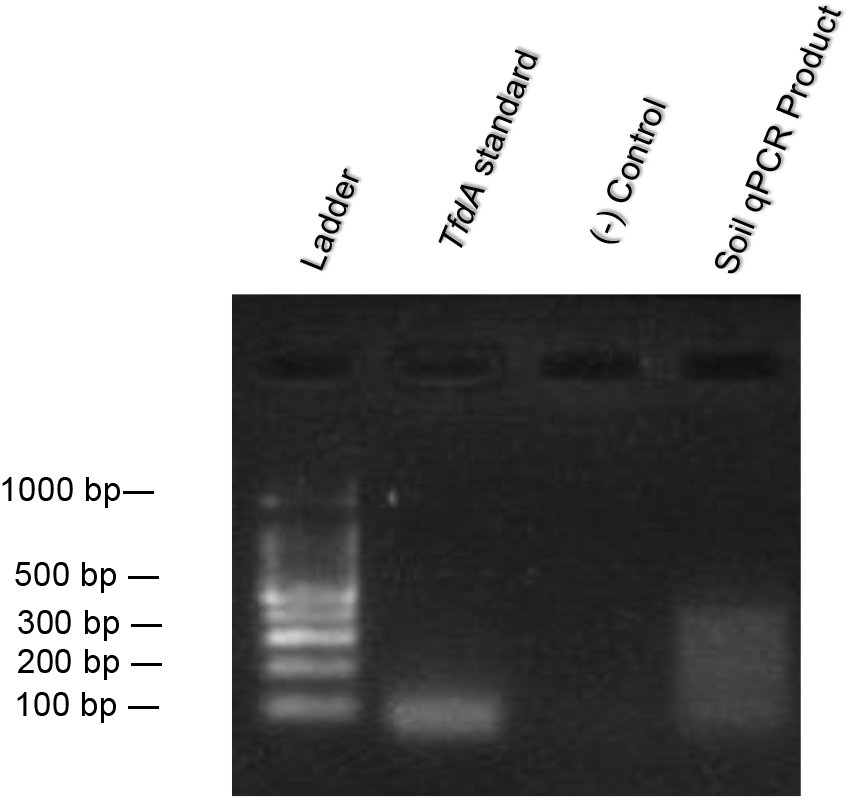
Electrophoresis gel of a qPCR amplification product of *tfdA* class I 81 bp primer in soil sample with 100 bp ladder.

**Figure 2.**
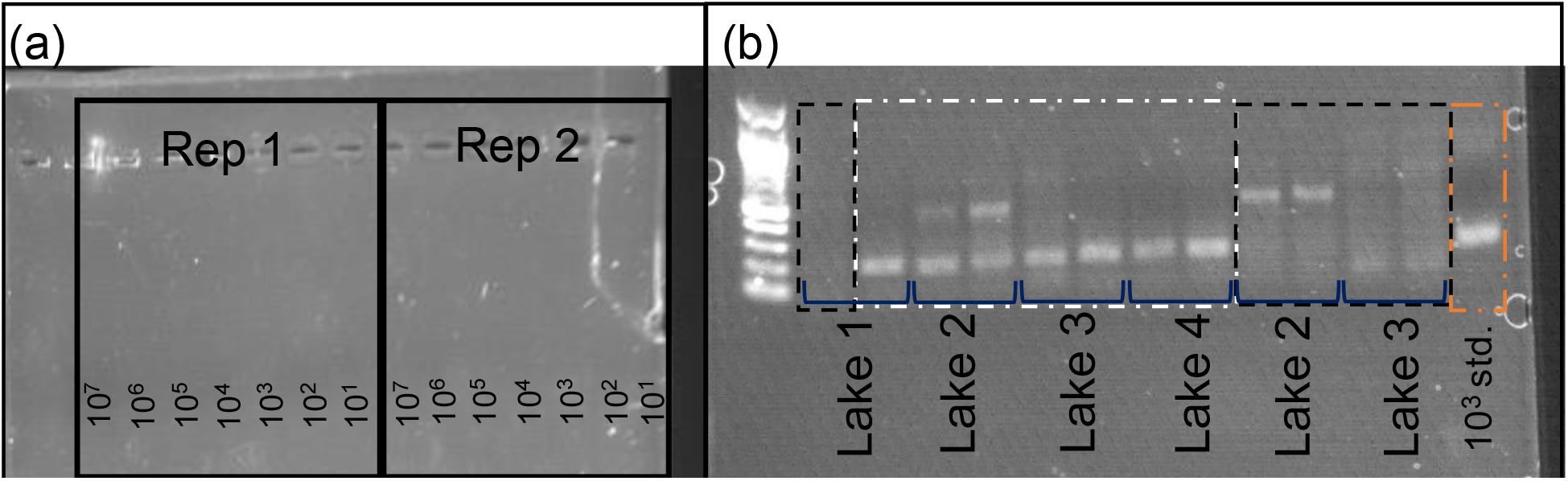
Gel of lake sediment recovery experiment using a serial dilution of *tfdA* class I gene standard from 10^7^ (most left) to 10^1^ (most right) (a). Gel of lake sediments and spiked with 10^3^ gene copies (white boxes) and unspiked sediments (black boxes) as well as a 10^3^ standard (orange box) (b). Ladder in both (a) and (b) is New England biotech 100 bp ladder. Band size is approximately 215 bp.

To evaluate interference by impurities co-extracted from sediments, serial dilutions of the class I *tfdA* gene from 10^7^ to 10^1^ copies were spiked into duplicate 0.5 g wet sediment aliquots previously unexposed to 2,4-D, extracted, and amplified with the 215 bp primer. High spiked gene copies (10^5^ and 10^2.5^ after correcting concentration for saturated sediments) had a percent recovery of 96-107% (**Table 2**), indicating successful recovery by DNA extraction and amplification of the standard. However, poor percent recoveries were observed at low copies (below 10^2.5^). Gene products visualized using a 1.5% v/v agarose gel showed some smearing and additional bands for all products (**Figure 2a**). At high concentrations, a prominent band was present at around 200 bp, which aligns with the expected fragment size of 215 bp. However, the band was less clear at 10^2.5^ copies and below, which also had poor calculated recoveries. This suggests, even with the standard derived from cultured representatives, low concentrations of the *tfdA* gene were not selectively amplified using the primers when the template DNA was present at relatively low abundance.

**Table 2.** Recovery of *tfdA* class I gene in sediments unexposed to 2,4-D. Expected gene recoveries are calculated based on sediment mass and water content.

| **Log10 gene**<br>**copies**<br>**Expected** | **Log10 gene**<br>**copies**<br>**Rep 1** | **Log10 gene**<br>**copies**<br>**Rep 2** | **Rep 1**<br>**recovery** | **Rep 2**<br>**recovery** | **Average**<br>**recovery** | **SD**<br>**recovery** |
| --- | --- | --- | --- | --- | --- | --- |
| 5.73 | 5.64 | 5.47 | 0.98 | 0.95 | 0.97 | 0.02 |
| 4.40 | 4.73 | 4.66 | 1.08 | 1.06 | 1.07 | 0.01 |
| 3.54 | 3.67 | 3.62 | 1.04 | 1.02 | 1.03 | 0.01 |
| 2.51 | 2.66 | 2.74 | 1.06 | 1.09 | 1.08 | 0.02 |
| 1.42 | 2.02 | 1.94 | 1.42 | 1.36 | 1.39 | 0.04 |
| -0.565 | 1.49 | 1.73 | -2.69 | -3.06 | -2.85 | 0.30 |
| -0.229 | 1.63 | 1.54 | -7.10 | -6.71 | -6.90 | 0.28 |
| 0.000 | 1.59 | 1.65 |  |  |  |  |

We conducted additional recovery experiments with sediment from four additional lakes recently unexposed to 2,4-D (4). We added 10^3^ copies of the *tfdA* gene and amplified with the 215 bp primer (**Figure 2b**). Sediments from both lakes with the added standard had a clear product band at ~200 bp, but the unspiked sediments had no clear band at 200 bp. Rather, these unspiked sediments had some smearing and light bands at 500 bp. This confirmed the finding that the primers could amplify the standard when found at relatively high levels in the extracted DNA and suggests that the lack of native *tfdA* amplification was not due to contaminants in the extraction interfering with the qPCR assay. Additionally, another non-specific product was being amplified.

Additional attempts to optimize the reaction did not improve amplification results. First, we attempted to adjust annealing temperatures by running a temperature gradient from 60 - 70°C and separately changing annealing time from 15 seconds to 30 seconds. We also tested doubling and tripling primer concentrations as well as lowering template concentrations to 0.5x and 0.25x of the standard protocol. Lastly, we performed additional clean up steps with phenol-chloroform and salt/alcohol precipitation to remove additional interferences prior to amplification. None of these troubleshooting steps generated clean bands on a gel from environmental samples. As a result, we cloned and sequenced PCR products to determine what was being amplified.

### Failed amplicon cloning and sequencing of PCR products

We constructed clone libraries to sequence individual amplicons from two soil (81 bp primer), one water (215 bp primer), and one sediment (215 bp primer) sample (**Table 1**). We selected 23 colonies for sequencing (5 from soil, 13 from water, 5 from sediment; **Table 1**). Following plasmid prep and amplification using M13 primers, visualization of 19 plasmid insert-derived PCR products from the sediment and water samples generated bands of approximately 200 bp each, which is shorter than the expected product of 215 bp. Amplified plasmid inserts from the remaining four clones from soil samples showed a band size between 200 and 500 bp (**Figure 3**). Analysis with blastn versus the NCBI nr database confirmed the sequences from samples A-S (water, sediment, and soil derived colonies) were fragments of the cloning vector rather than the *tfdA* gene product. For samples T-W (soil only) the insert sequence had less than 20% alignment with representative sequences from Class I, II, and III *tfdA* genes (**Table S1**). Sequences producing significant alignments to the insert sequence were not homologous to TfdA.

**Figure 3.**
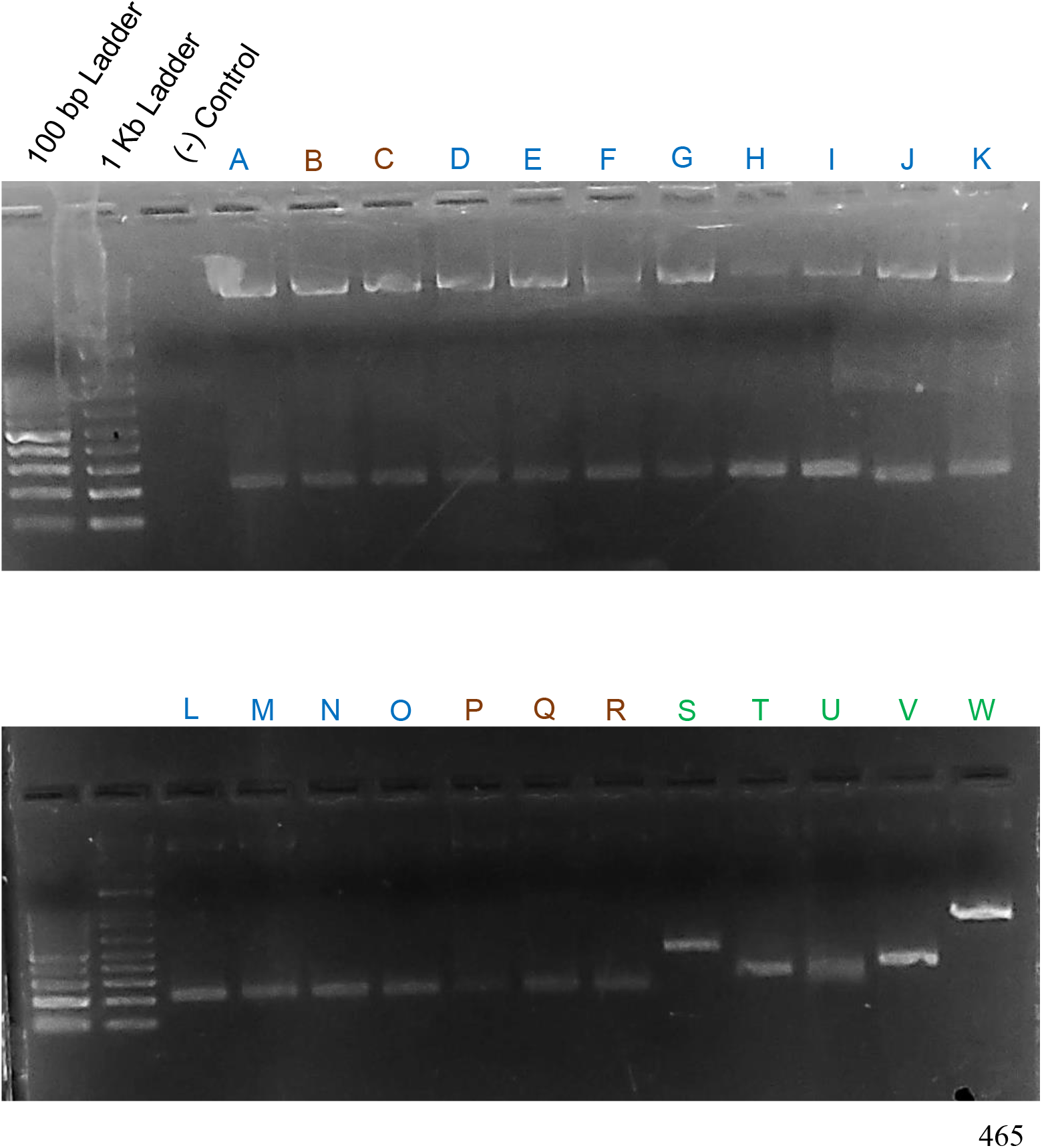
Gel of colonies with gene product from two soil (green letters), one sediment (brown letters), and one water sample (blue letters) after TOPO cloning, incubation in LB and plates at 37°C, and amplified using M13 primers. Expected product size of 215 bp for samples A-R and 81 bp for samples S-W. Ladder size indicated on gel is 100 bp and 1 kbp from left to right.

Our work here shows the *tfdA* qPCR primers failed to detect 2,4-D degrading microorganisms in environmental samples, which is consistent with other studies that have demonstrated challenges in applying primers created from cultured representatives to environmental samples (36–39). A potential explanation for the failed amplification results despite the observed degradation of 2,4-D in environmental samples could be the presence of the *cad* pathway, which can degrade 2,4-D and has predominantly been found in *Burkholderia* (40–42). Additionally, the isolation technique used can influence what class of *tfdA* is recovered, suggesting primers created from isolates may miss parts of the degrading community solely based on how the strains were isolated (43). Thus, we further investigated if extensive *tfdA* sequence divergence at the primer binding site could explain why the primers failed on our environmental samples using non-isolation-based tools.

### Phylogenetic analysis leads to discovery of putative degraders

We searched for *tfdA* homologs using a custom Hidden Markov Model (HMM) in more than 200,000 publicly available isolate genomes and MAGs available in Genbank at the time of the search and found 1,035 putative 2,4-D degraders spanning over a dozen phyla and several different environmental sources (**Figure 4**, **Metadata file available on <u>FigShare Project Page</u>)**. 90% of identified putative degraders belong to *Proteobacteria*, specifically the *Alpha-, Beta-, Gamma-, Deltaproteobacteria* classes and otherwise unresolved Proteobacteria (**Figure 4b, Figure 4c**), groups which have previously been identified to contain 2,4-D degraders. The remaining 103 novel putative degraders are classified within the Acidobacteria (n = 1), Actinobacteria (n = 5), Fibrobacteres (n = 1), Gemmatimonadetes (n = 2), and PVC Superphylum (n = 7), as well as several recently described candidate phyla, including *Candidatus* Eremiobacteraeota (n = 20), *Candidatus* Latescibacteria (n = 1), *Candidatus* Rokubacteria (n = 61), and *Candidatus* Tectomicrobia (n = 4). Putative degrader genomes were recovered from marine, freshwater, and wetland aquatic environments in addition to the more well studied terrestrial and subsurface environments. The distribution of the putative degraders across all environments and phyla aligns with evidence for horizontal gene transfer (28). Importantly, cultured and experimentally verified representatives clustered separately from the putative degraders (**Figure 4a**), suggesting the gene is highly conserved among degraders that are easily isolated or can use 2,4-D as a sole carbon source. This also suggests that the three previously described classes of *tfdA* represent a small portion of the overall diversity of *tfdA*.

**Figure 4.**
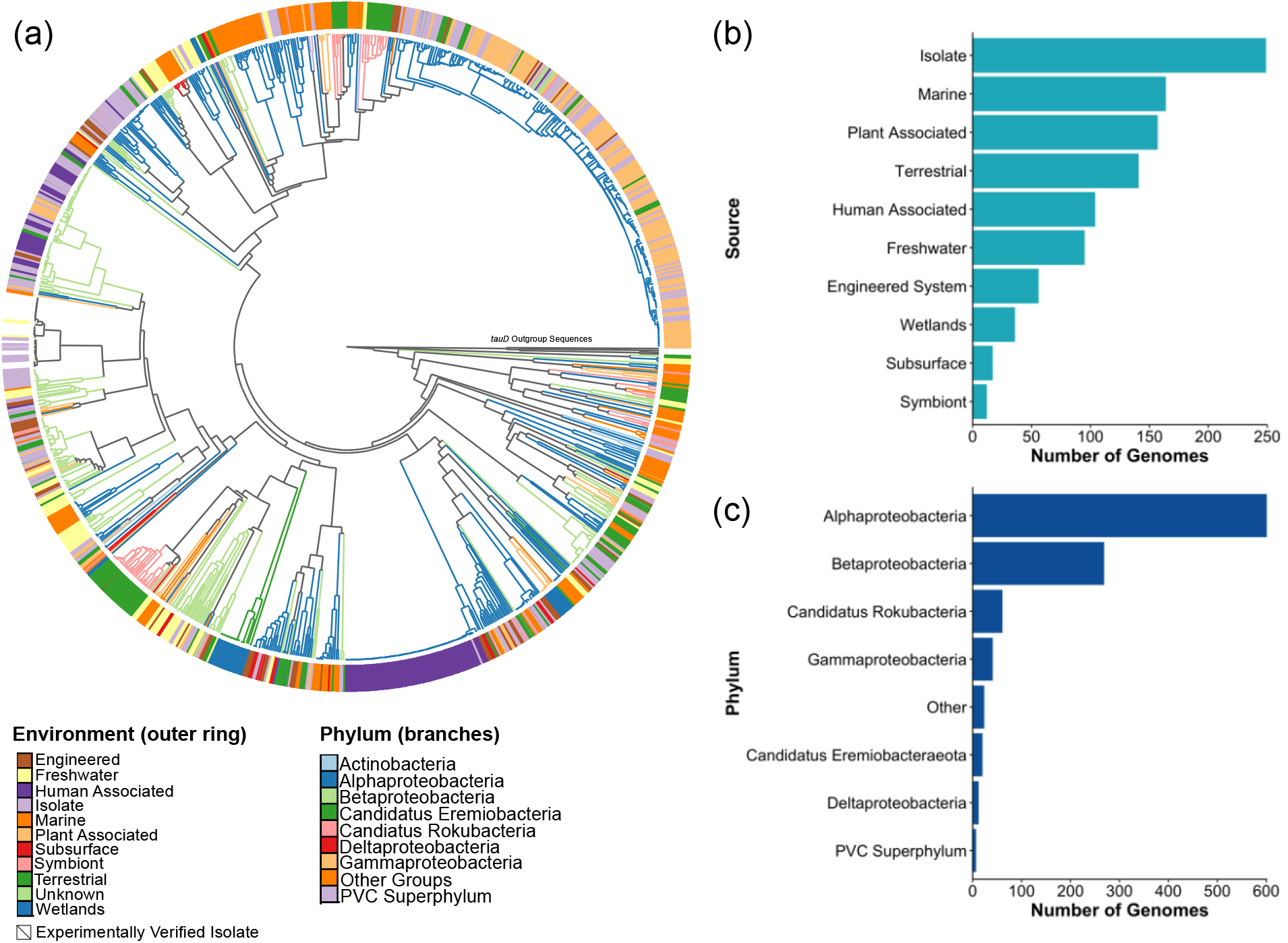
Phylogenetic tree of putative *tfdA* degraders (a). Tree color corresponds to phylum and outer ring corresponds to environmental origin. Counts of sequences from each environment of origin of putative 2,4-D degraders (b). Counts of sequences from each phyla of putative 2,4-D degraders (c).

We found putative 2,4-D degraders in Acidobacteria, Gemmatimonadetes, and the PVC superphylum that have been previously reported as degraders or members of the degrading community of the related phenoxyalkanoic acid herbicide (2-methyl-4-chlorophenoxy) propionic acid (MCPA) (44). While these phyla have not been shown to degrade 2,4-D, the structural similarity of MCPA to 2,4-D and the demonstrated ability of *tfdA* to degrade both herbicides (29, 30, 45) suggests these microbes have the potential to degrade 2,4-D as well.

Most of the novel putative degraders were members of *Candidatus* Rokubacteria in our search and contained representatives from marine, freshwater, soil, subsurface, and sediment environments. This observation is consistent with previous reports of the global diversity of Rokubacteria (46), which are also expected to play an important role in sulfur cycling (47, 48) and degradation of complex carbon compounds found in leaf litter or root exudates (48–50). Previous work demonstrating uncultured microbes like Rokubacteria may use alternative metabolic pathways that prevent their culturing under typical conditions (46, 51, 52) highlights the power of non-culture-based methods in exploring community structure and function and the limits of historic cultured-based primer development methods for environmental analyses.

Wetland-derived putative degraders are predominantly from *Candidatus* Eremiobacteraeota, specifically found in artic peatlands. Another uncultured phylum, Eremiobacteraeota are described as “heterotrophic scavengers” (53) found in polar/alpine environments (54–57), as well as polychlorinated biphenyl- and fracking-contaminated environments (58, 59). All instances of *Candidatus* Eremiobacteraeota in our study were derived from the arctic wetland environment. Other wetland-derived putative degraders include *Acetobacteraceae*, *Rhizobiales*, and *Rhodospirillales* (Alphaproteobacteria), *Burkholderiales* (Betaproteobacteria), nine unresolved Gammaproteobacteria, and one unresolved Deltaproteobacteria.

Several of the putative degraders from all environments have been found in heavily contaminated environments. For example, *Sphingomonadaceae* (Alphaproteobacteria) are known to have several members that can tolerate or degrade contaminants, including 2,4-D (25, 28, 60, 61), and have been found in gasoline (62), polyaromatic hydrocarbon (63), hexachlorocyclohexane (64), copper mine (65), and electronic waste (66) contaminated soils and sediments. We found *Sphingomonadaceae* in all sampled environments, including a large group of human-associated isolates from building infrastructure and patient cultures during an investigation of an outbreak of multiple-drug resistant *S. koreensis* at the NIH Clinical Center from 2006-2016 (67). We found several other *tfdA* carrying representatives in drinking water samples, specifically a member of the family *Bradyrhizobiaceae* (68, 69), *Variovorax paradoxus* (70), and other Proteobacteria representatives (69). Additional putative 2,4-D degraders known to tolerate contaminants include the isoprene-degrading *Variovorax* (71), heavy metal-resistant *Altererythrobacter atlanticus* (72), and the previously described *Eremiobacteraeota* (58, 59).

The variety in origin and diversity of putative 2,4-D degraders suggests they can be found nearly ubiquitously across natural and built environments and supports previous evidence that the gene can be horizontally transferred. Many of the degraders identified here, both established and putative, are associated with complex carbon cycling, such as the Eremiobacteraeota and Rokubacteria. This finding aligns with previous evidence that *tfd* genes or ancestral *tfd* genes were used to degrade naturally occurring compounds that are structurally similar to 2,4-D (73–75). Additionally, the large number of putative degraders found in contaminated or engineered environments implies the gene or plasmid is resilient or conveys advantageous traits in stressful environments, even if 2,4-D or MCPA is not present.

### Failed *in silico* amplification with qPCR primers on putative degraders

Given the wide distribution of the *tfdA* gene across all environments sampled here, we investigated primer accuracy by comparing *in silico* amplification using the 81 and 215 bp primers from putative degrader genomes with confirmed 2,4-D degrader sequences. Applying the 81 and 215 bp primers to the curated sequences of putative degraders resulted in poor rates of *in silico* amplification, even when allowing for one and three mismatches (**Figure 5a**). The 81 bp forward and reverse primer amplified zero sequences out of 1604 sequences when allowing for zero mismatches, 10 sequences when allowing for one mismatch, and 55 sequences when allowing for three mismatches (i.e., 3.4% success rate). The 215 bp primer had slightly better success with 8 amplified sequences assuming zero mismatches. Increasing the number of allowable mismatches to one and three resulted in 9 and 83 (i.e., maximum 5.2% success rate) amplified sequences, respectively (**Figure 5a**).

**Figure 5.**
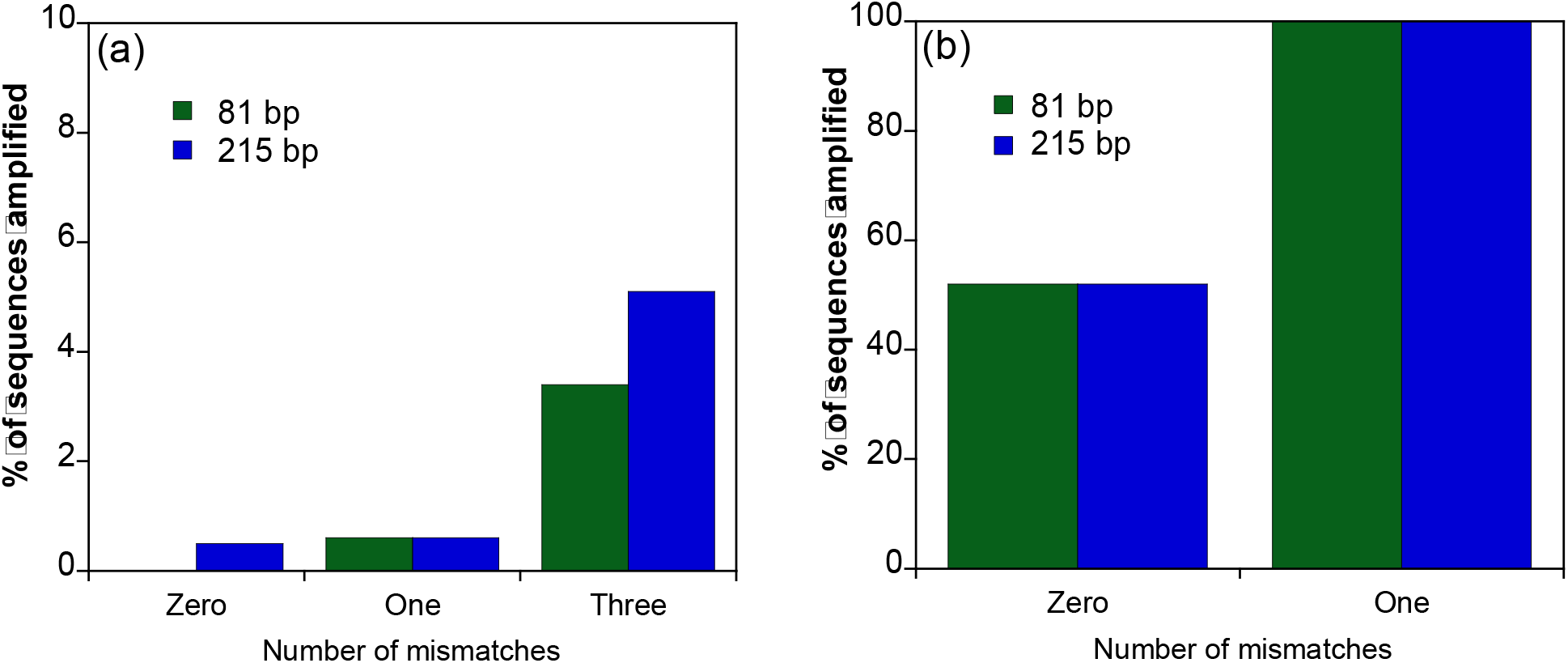
Amplification success rates using 1604 sequences of putative 2,4-D degraders tested with 81 and 215 bp primers (a). Zero, one, and three mismatches were allowed. Amplification success rates of 29 experimentally verified 2,4-D degraders with 81 and 215 bp primers (b).

In contrast, applying both primer sets to experimentally verified and cultured 2,4-D degraders (25) resulted in 52% amplification with zero mismatches and 100% amplification for one mismatch (**Figure 5b**). These results, combined with the phylogenetic tree, show the diversity of the *tfdA* gene is much larger than previously described using when using *tfdA* sequences obtained from microorganisms predominantly cultivated from soil environments.

## Conclusion

In this study we expanded the diversity of known putative 2,4-D degraders using genome-resolved metagenomics and evaluated the accuracy of published qPCR primers commonly used to quantify the amount of *tfdA* gene in environmental samples. Our study found a significant gap between amplification of experimentally verified degraders and putative environmental degraders, which demonstrates how the use of cultured representatives to develop quantitative molecular tools has been known to underestimate the diversity of gene-carrying microorganisms in natural environments (51). Additionally, many of the putative degrader sequences were within phyla or families containing experimentally verified degraders, groups known to degrade similar phenoxyacid herbicides, or groups known to degrade other organic contaminants across the natural and built environment, suggesting the gene is more widespread than previously described.

Characterizing the *tfdA* gene distribution and prevalence across the natural environment is critical for understanding herbicide biodegradation, which is especially important for relatively persistent compounds such as 2,4-D. Our study evaluated established qPCR primers for the *tfdA* gene through molecular and computational techniques and found these primers missed a majority of the potential 2,4-D degrading community. We did not evaluate PCR primers that predated the development of the qPCR primers (34, 76), nor did we explore the more recently reported *cad* genes (41). Our results underscore the importance of using caution when applying qPCR primers developed from a small number of sequences from cultured representatives to environmental samples because they are likely to misrepresent true environmental diversity. Additionally, further investigation into the 2,4-D degrading community is critical to understand the true environmental impact of phenoxyacid herbicides or other persistent halogenated compounds that are prevalent in the environment. The combination of targeted molecular techniques and genome-resolved metagenomics is a powerful way to link quantitative data to qualitative community and function data for a holistic understanding of microbial community response to anthropogenic stressors and both should be considered when designing future studies.

## Methods

### Environmental sample collection

Lake water and sediment were collected from several Wisconsin Lakes undergoing large-scale 2,4-D treatments (i.e., lake wide concentrations >0.45 μM), described in detail in (4). Water and sediment used in TOPO cloning were from Random Lake, which had whole-lake 2,4-D treatments in 2018 and 2019. The water column biomass and sediment used in TOPO cloning were collected May 20 and 29, 2019, respectively. These samples were chosen for additional analysis because of a low Ct-value (i.e., high initial gene copies) relative to other samples in preliminary qPCR tests. Water samples were collected via grab sampling from the epilimnion of the deepest point of the lake (< 1 m sampling depth) and stored on ice until processing. 1.5 L of water was filtered through a 0.22 μm polyethersulfone filter in a Sterivex filter unit (Millipore Sigma) and stored in a −20°C freezer. Lake sediment samples were collected by hand using a 5 cm diameter PVC core tube. Three individual sediment cores (top 0-4 cm) were homogenized in a sterile plastic bag and transferred to a 50 mL falcon tube for long term storage in a −20°C freezer. Soil samples were collected using a 100 mm PVC diameter soil core at a depth of 200 mm from OJ Noer Turfgrass Research Station in Madison, Wisconsin, on July 30, 2020. 2,4-D formulation was applied to field plots using a nozzle pressure of 40 psi using a CO_2_ pressurized boom sprayer with two XR Teejet AI8004 nozzles. The herbicide was agitated by hand and applied at a rate equivalent to 0.35 mL / m^2^ of the commercial 2,4-D product with an initial concentration of 11.9 μM. Soil subsamples from each core were then collected in the following regions: upper soil (top 5 cm) and lower soil (15-20 cm depth) using a stainless-steel soil core sampler.

### qPCR amplification

Sediment, water, and soil samples exposed to 2,4-D were collected and amplified as described previously in (4) and (77). DNA was extracted using an MP Bio Fast DNA Spin Kit and quantified for DNA concentration using an Invitrogen Qubit 3.0 fluorimeter. Amplification was attempted using non-class specific *tfdA* primers intended to amplify an 81bp or 215bp fragment (29, 30).

Primer sequences and thermocycle conditions were adapted from Baelum et al. 2008 (30). For the 215 bp product, the forward primer was 5’-GAGCACTACGCRCTGAAYTCCCG-3’ and the reverse primer was 5’-GTCGCGTGCTCGAGAAG-3’. Amplification was done using a BioRad Thermocycler with initial heating for 10 minutes at 95°C and 40 cycles of 15 seconds at 95°C; 30 seconds at 64°C, and 30 seconds at 72°C. For the 81 bp product, forward primer was 5’-GAGCACTACGCRCTGAAYTCCCG-3’ and the reverse primer was 5’-SACCGGMGGCATSGCATT-3’. The reaction conditions were as follows: 3 min at 95°C for enzyme activation, followed by 35 cycles of 15 s at 95°C and 1 min at 62°C in a Bio-Rad CFX96 Touch Real-Time PCR Detection System. Amplified product melt curves were inspected to assess product homogeneity and visualized using agarose gel electrophoresis (1-1.5% wt/vol) to confirm product length.

### Bacterial strains and *tfdA* reference genes

*tfdA* class I gene, originally from *R. eutropha* pJP4(JMP134), *tfdA* Class II and Class III genes originated from *Burkholderia* strain RASC and *Burkholderia cepacia* strain 2a pIJB1, respectively, were synthesized by Integrated DNA Technologies in a pUCIDT- AMP cloning vector and transformed into *E. coli* AMP-resistant competent cells and processed as described in (77). Cultures of each class were grown overnight at 37°C on Luria Bertani (LB) medium containing 500 mg/L ampicillin. According to the manufacturer’s protocol, DNA plasmid isolations were performed using E.Z.N.A. Plasmid DNA Kit (Omega Bio Tek, Radnor, PA). Plasmid DNA was quantified using a NanoDrop ND-1000 spectrophotometer (ThermoFisher Scientific, Waltham, MA). Plasmids were serially diluted and used as standards for quantitative real-time PCR (29).

### TOPO cloning

TOPO cloning using an Invitrogen TOPO^®^-TAcloning (Invitrogen, Karlruhe, Germany) kit with One Shot TOP10 chemically competent *E. coli* cells was conducted on two soil, one sediment, and one water sample amplified with either the 81 bp or 215 bp primer set. To set up the TOPO^®^ reaction, 4 μL of fresh qPCR product, 1 μL salt solution and 1 μL of the pCR™^4^-TOPO^®^ vector were combined. Fresh PCR product was inserted into the provided plasmids and transformed into the chemically competent cells. Cells were spread on LB Agar plates with 50 mg/mL kanamycin, incubated overnight at 37°C. Colonies grown on overnight plates were transferred to LB broth with 50 mg/mL kanamycin and incubated at 37°C overnight again. Aliquots of liquid cultures were extracted using an Invitrogen PureLink Quick Plasmid Miniprep kit. Plasmids were then amplified with IDT ReadyMade M13 (−20) forward primers. Product sizes were evaluated using gel electrophoresis (1.5% agarose) with SYBR Safe DNA Gel Stain (Fisher Scientific, Chicago, IL) and purified using a Qiagen PCR Purification Kit (Qiagen, Hilden, Germany). Final products were quantified using a NanoDrop 1000 (Thermo Fisher Scientific, Waltham, MA) and sequenced on a 3730XL Genetic Analyzer (ThermoFisher) for Sanger sequencing in the Biotechnology Center at the University of Wisconsin-Madison. Obtained sequences were processed using A Plasmid Editor (ApE) (78) and analyzed using NCBI NucleotideBLAST (blastn) and blastx.

### Accessed datasets

We used a combination of publicly available sequenced isolates and metagenome-assembled genomes (MAGs) to survey *tfdA* diversity. All publicly available isolates and MAGs were accessed from Genbank in August 2019, resulting in more than 200,000 genomes to search for *tfdA*. We also included MAGs assembled from three freshwater lakes, Lake Tanganyika in the East African Rift Valley (79), Trout Bog near Minocqua, WI, and Lake Mendota in Madison, WI (80). Samples from Trout Bog and Lake Mendota were sequenced and population genomes assembled as described previously (80). Samples were sequenced and population genomes were also assembled from two stations in Lake Tanganyika (i.e., Kegoma and Mahale) as described previously (79).

### Identification of putative 2,4-D degraders

We constructed a Hidden Markov Model (HMM) of the TfdA protein using a collection of 30 reference TfdA protein sequences from experimentally verified isolates with 2,4-D degradation activity (25). Reference protein sequences were aligned with MUSCLE and an HMM built with the hmmbuild function of the HMMER suite (81). The constructed HMM profile was used to then identify putative 2,4-D degraders among over 1000 publicly available genomes, including pure culture isolates and metagenome assembled genomes (MAGs). For all genome sequences, open reading frames and protein-coding genes were predicted with Prodigal (82). We used hmmsearch as part of the HMMER program to search all predicted proteins for putative TfdA sequences with an e-value cutoff of 1e-50 (81). All sequence hits were aligned with MUSCLE and visualized using AliView to manually check for specific conserved residues (83, 84). Based on the predicted positions of active site residues of TfdA in *Cupriavidus pinatubonensis* JMP134 (previously known as *Cupriavidus necator* JMP134 and *Ralstonia eutropha* JMP134), we checked for the presence of residues at His263 and Arg278 that appeared to be required for degradation and mostly conserved among the majority of sequences (75, 85, 86). Any sequences without these residues aligned with the corresponding positions of *C. pinatubonensis* were discarded.

### Phylogenetic diversity of putative 2,4-D degraders functional annotations

A phylogenetic tree of all TfdA protein sequences was created from the alignment of all confident TfdA hits and constructed with RAxML (87). The tree was rooted using TauD sequences as an outgroup, as these sequences are from a related family of dioxygenases as TfdA (61, 88, 89). TauD sequences from *Mycobacterium marinum* strain M (ACC39598), *Burkholderia pseudomallei* 1710b (ABA48168), *Escherichia coli* K12 (BAE76149.1), and *Yersinia pestis* CO92 (CAL18870.1) were used as outgroup dioxygenases, as these were also used in Gonod et al. 2006 (88). These outgroups sequences were used to root the tree, and any putative TfdA sequences that did not fall within a monophyletic clade outside of the root were removed. The environmental or isolation source was overlaid on the tree as described by the project type for the associated BioProject for each strain or MAG in Genbank. The tree and associated metadata were visualized using EMPress v1.1.0 (90), and a full phylogenetic tree with bootstrap values is provided in the Supplementary Data.

### *In silico* primer analysis

Primer specificity was evaluated using the Geneious (Biomatters, New Zealand) primer mapper. Both the 215 and 81 bp forward and reverse primer sets were mapped against a database of *tfdA* sequences from cultured organisms and environmental MAGs. Mapping was done with different stringency criteria, allowing for 0, 1, and 3 mismatches. Sequences that were not matched by the primers when three mismatches were allowed were considered unlikely to be amplified by the primers in environmental samples.

## Data availability statement

All data files and supplementary information are available at https://figshare.com/projects/Expanded_diversity_of_tfdA_harboring_bacteria_across_the_natural_and_built_environment/145275. Metadata, HMM, phylogenetic tree with bootstraps, and alignment files for curated sequences well as Sanger sequencing results for 215 bp primer verification and TOPO cloning are available. A PDF Supplementary File with Table S1 and Figure S1 is also available. All code available at https://github.com/elizabethmcd/tfdA.

## Acknowledgements

Thank you to Dr. Jan Roelof van der Meer from the University of Lausanne, Switzerland for supplying cultures for creation of the qPCR standards. We acknowledge valuable input from Angela Magness and Naila Barbosa da Costa in the qPCR and sequencing for this project. This research was performed in part using the Wisconsin Energy Institute computing cluster, which is supported by the Great Lakes Bioenergy Research Center as part of the U.S. Department of Energy Office of Science (DE-SC0018409), as well as the Center for High-Throughput Computing (CHTC) at UW-Madison in the Department of Computer Sciences. CHTC is supported by UW-Madison, the Advanced Computing Initiative, the Wisconsin Alumni Research Foundation, the Wisconsin Institute for Discovery, and the National Science Foundation. Additional funding provided by the National Institute of Food and Agriculture, United States Department of Agriculture Hatch/Evans-Allen/McIntire Stennis project #1013114, Wisconsin Department of Natural Resources, National Science Foundation Graduate Research Fellowship (A.M.W. and B.D.P), UW-Madison Department of Bacteriology Predoctoral Fellowship (E.A.M), an O. N. Allen Small Grant (A.G.V), and an Anna Grant Birge Memorial Award (A.M.W.).

## Notes

https://figshare.com/projects/Expanded_diversity_of_tfdA_harboring_bacteria_across_the_natural_and_built_environment/145275.

### Competing Interest Statement

The authors have declared no competing interest.

## References

1. Peterson MA, McMaster SA, Riechers DE, Skelton J, Stahlman PW. 2016. 2,4-D past, present, and future: A review. Weed Tech 30:303–345.

2. Nault ME, Netherland MD, Mikulyuk A, Skogerboe JG, Asplund T, Hauxwell J, Toshner P. 2014. Efficacy, selectivity, and herbicide concentrations following a whole-lake 2,4-D application targeting Eurasian watermilfoil in two adjacent northern Wisconsin lakes. Lake and Resvr Mange 30:1–10.

3. Góngora-Echeverría VR, Martin-Laurent F, Quintal-Franco C, Lorenzo-Flores A, Giácoman-Vallejos G, Ponce-Caballero C. 2019. Dissipation and adsorption of 2,4-D, atrazine, diazinon, and glyphosate in an agricultural soil from Yucatan State, Mexico. Water, Air, Soil Poll 230:131.

4. White AM, Nault ME, McMahon KD, Remucal CK. 2022. Synthesizing laboratory and field experiments to quantify dominant transformation mechanisms of 2,4-dichlorophenoxyacetic acid (2,4-D) in aquatic environments. Environ Sci Technol 56:10838–10848.

5. Dehnert GK, Freitas MB, DeQuattro ZA, Barry T, Karasov WH. 2018. Effects of low, subchronic exposure of 2,4-dichlorophenoxyacetic acid (2,4-D) and commercial 2,4-D formulations on early life stages of fathead minnows (Pimephales promelas). Envr Tox and Chem 37:2550–2559.

6. Dehnert GK, Freitas MB, Sharma PP, Barry TP, Karasov WH. 2021. Impacts of subchronic exposure to a commercial 2,4-D herbicide on developmental stages of multiple freshwater fish species. Chemosphere 263:127638.

7. Chavez Rodriguez L, Ingalls B, Schwarz E, Streck T, Uksa M, Pagel H. 2020. Gene-centric model approaches for accurate prediction of pesticide biodegradation in soils. Environ Sci Technol 54:13638–13650.

8. Ogram A v., Jessup RE, Ou LT, Rao PSC. 1985. Effects of sorption on biological degradation rates of (2,4-dichlorophenoxy)acetic acid in soils. Appl Environ Microbiol 49:582–587.

9. Han L, Zhao D, Li C. 2015. Isolation and 2,4-D-degrading characteristics of *Cupriavidus campinensis* BJ71. Braz J Microbiol 46:433–441.

10. Zabaloy MC, Gómez MA. 2014. Isolation and characterization of indigenous 2,4-D herbicide degrading bacteria from an agricultural soil in proximity of Sauce Grande River, Argentina. Annals Micro 64:969–974.

11. Paszko T, Muszyński P, Materska M, Bojanowska M, Kostecka M, Jackowska I. 2016. Adsorption and degradation of phenoxyalkanoic acid herbicides in soils: A review. Envr Tox and Chem 35:271–286.

12. Fukumori F, Hausinger RP. 1993. *Alcaligenes eutrophus* JMP134 “2,4-dichlorophenoxyacetate monooxygenase” is an α-ketoglutarate-dependent dioxygenase. J Bact 175:2083–2086.

13. Kumar A, Trefault N, Olaniran AO. 2016. Microbial degradation of 2,4-dichlorophenoxyacetic acid: Insight into the enzymes and catabolic genes involved, their regulation and biotechnological implications. Crit Rev Micro 42:194–208.

14. Laemmli CM, Leveau JHJ, Zehnder AJB, Roelof Van Der Meer J. 2000. Characterization of a second *tfd* gene cluster for chlorophenol and chlorocatechol metabolism on plasmid pJP4 in *Ralstonia eutropha* JMP134 (pJP4). J Bact 182:4165–4172.

15. Zuanazzi NR, Ghisi N de C, Oliveira EC. 2020. Analysis of global trends and gaps for studies about 2,4-D herbicide toxicity: A scientometric review. Chemosphere 241:125016.

16. Liu W, Li H, Tao F, Li S, Tian Z, Xie H. 2013. Formation and contamination of PCDD/Fs, PCBs, PeCBz, HxCBz and polychlorophenols in the production of 2,4-D products. Chemosphere 92:304–308.

17. Islam F, Wang J, Farooq MA, Khan MSS, Xu L, Zhu J, Zhao M, Muños S, Li QX, Zhou W. 2018. Potential impact of the herbicide 2,4-dichlorophenoxyacetic acid on human and ecosystems. Environ Int 111:332–351.

18. Atwood D, Paisley-Jones C. 2017. US EPA - Pesticides Industry Sales and Usage 2008 - 2012. https://www.epa.gov/sites/default/files/2017-01/documents/pesticides-industry-sales-usage-2016_0.pdf. Accessed 2020-08-10.

19. Barbosa AMC, Solano M de LM, Umbuzeiro G de A. 2015. Pesticides in drinking water – The Brazilian monitoring program. Front Public Health 3:246.

20. Nicolaisen MH, Bælum J, Jacobsen CS, Sørensen J. 2008. Transcription dynamics of the functional *tfdA* gene during MCPA herbicide degradation by *Cupriavidus necator* AEO106 (pRO101) in agricultural soil. Environ Microbiol 10:571–579.

21. Chaudhry GR, Huang GH. 1988. Isolation and characterization of a new plasmid from a *Flavobacterium* sp. which carries the genes for degradation of 2,4-dichlorophenoxyacetate. J Bact 170:3897–3902.

22. Suwa Y, Wright AD, Fukimori F, Nummy KA, Hausinger RP, Holben WE, Forney LJ. 1996. Characterization of a chromosomally encoded 2,4-dichlorophenoxyacetic Acid/ α-Ketoglutarate dioxygenase from *Burkholderia* sp. strain RASC. Appl Environ Microbiol 62:2464–2469.

23. Matheson VG, Forney LJ, Suwa Y, Nakatsu CH, Sexstone AJ, Holben WE. 1996. Evidence for acquisition in nature of a chromosomal 2,4-dichlorophenoxyacetic acid/ α-ketoglutarate dioxygenase gene by different *Burkholderia* spp. Appl Environ Microbiol 62:2457–2463.

24. Lerch TZ, Dignac MF, Barriuso E, Bardoux G, Mariotti A. 2007. Tracing 2,4-D metabolism in *Cupriavidus necator* JMP134 with ^13^C-labelling technique and fatty acid profiling. J Micro Methods 71:162–174.

25. Baelum J, Jacobsen CS, Holben WE. 2010. Comparison of 16S rRNA gene phylogeny and functional *tfdA* gene distribution in thirty-one different 2, 4-dichlorophenoxyacetic acid and 4-chloro-2-methylphenoxyacetic acid degraders. Syst Appl Microbiol 33:67–70.

26. Marron-Montiel E, Ruiz-Ordaz N, Rubio-Graados C, Juarez-Ramirez C, Galindez-Mayer CJ. 2006. Isolation, kinetic characterization in batch and continuous culture and application for bioaugmenting an activated sludge microbial community. Process Biochem 41:1521–1528.

27. Musarrat J, Bano N. 2000. Isolation and characterization of 2,4-dichlorophenoxyacetic acid-catabolizing bacteria and their biodegradation efficiency in soil. Word Jrnl Micro Biotech 16:495–497.

28. McGowan C, Fulthorpe R, Wright A, Tiedje JM. 1998. Evidence for interspecies gene transfer in the evolution of 2,4-dichlorophenoxyacetic acid degraders. Appl Environ Microbiol 64:4089–4092.

29. Bælum J, Jacobsen CS. 2009. TaqMan probe-based real-time PCR assay for detection and discrimination of class I, II, and III *tfdA* genes in soils treated with phenoxy acid herbicides. Appl Environ Microbiol 75:2969–2972.

30. Bælum J, Nicolaisen MH, Holben WE, Strobel BW, Sørensen J, Jacobsen CS. 2008. Direct analysis of *tfdA* gene expression by indigenous bacteria in phenoxy acid amended agricultural soil. ISME 2:677–687.

31. Batoǧlu-Pazarbaş M, Milosevic N, Malaguerra F, Binning PJ, Albrechtsen HJ, Bjerg PL, Aamand J. 2013. Discharge of landfill leachate to streambed sediments impacts the mineralization potential of phenoxy acid herbicides depending on the initial abundance of *tfdA* gene classes. Environ Pollut 176:275–283.

32. Batioǧlu-Pazarbaşi M, Bælum J, Johnsen AR, Sørensen SR, Albrechtsen HJ, Aamand J. 2012. Centimetre-scale vertical variability of phenoxy acid herbicide mineralization potential in aquifer sediment relates to the abundance of *tfdA* genes. FEMS Microbiol Ecol 80:331–341.

33. Stibal M, Bælum J, Holben WE, Sørensen SR, Jensen A, Jacobsen CS. 2012. Microbial degradation of 2,4-dichlorophenoxyacetic acid on the Greenland ice sheet. Appl Environ Microbiol 78:5070–5076.

34. de Lipthay JR, Aamand J, Barkay T. 2006. Expression of *tfdA* genes in aquatic microbial communities during acclimation to 2,4-dichlorophenoxyacetic acid. FEMS Microbiol Ecol 40:205–214.

35. de Lipthay JR, Tuxen N, Johnsen K, Hansen LH, Albrechtsen HJ, Bjerg PL, Aamand J. 2003. In situ exposure to low herbicide concentrations affects microbial population composition and catabolic gene frequency in an aerobic shallow aquifer. Appl Environ Microbiol 69:461–467.

36. Peterson BD, McDaniel EA, Schmidt AG, Lepak RF, Janssen SE, Tran PQ, Marick RA, Ogorek JM, DeWild JF, Krabbenhoft DP, McMahon KD. 2020. Mercury methylation genes identified across diverse anaerobic microbial guilds in a eutrophic sulfate-enriched lake. Environ Sci Technol 54:15840.

37. Gionfriddo CM, Wymore AM, Jones DS, Wilpiszeski RL, Lynes MM, Christensen GA, Soren A, Gilmour CC, Podar M, Elias DA. 2020. An improved *hgcAB* primer set and direct high-throughput sequencing expand Hg-methylator diversity in nature. Front Microbiol 11:541554.

38. Jones DS, Walker GM, Johnson NW, Mitchell CPJ, Coleman Wasik JK, Bailey JV. 2019. Molecular evidence for novel mercury methylating microorganisms in sulfate-impacted lakes. ISME 13:1659–1675.

39. Beach NK, Noguera DR. 2019. Design and assessment of species-level qPCR primers targeting comammox. Front Microbiol 10:1–15.

40. Itoh K, Kanda R, Sumita Y, Kim H, Kamagata Y, Suyama K, Yamamoto H, Hausinger RP, Tiedje JM. 2002. *tfdA*-like genes in 2,4-dichlorophenoxyacetic acid-degrading bacteria belonging to the *Bradyrhizobium-Agromonas-Nitrobacter-Afipia* cluster in α-*Proteobacteria*. Appl Environ Microbiol 68:3449–3454.

41. Kijima K, Mita H, Kawakami M, Amada K. 2018. Role of *CadC* and *CadD* in the 2,4-dichlorophenoxyacetic acid oxygenase system of *Sphingomonas agrestis* 58-1. J Biosci Bioengr 125:649–653.

42. Nielsen TK, Xu Z, Gözdereliler E, Aamand J, Hansen LH, Sørensen SR. 2013. Novel insight into the genetic context of the *cadAB* genes from a 4-chloro-2-methylphenoxyacetic acid-degrading *Sphingomonas*. PLoS One 8:e83346.

43. Lee TH, Kurata S, Nakatsu CH, Kamagata Y. 2005. Molecular analysis of bacterial community based on 16S rDNA and functional genes in activated sludge enriched with 2,4-dichlorophenoxyacetic acid (2,4-D) under different cultural conditions. Microb Ecol 49:151–162.

44. Liu YJ, Liu SJ, Drake HL, Horn MA. 2011. Alphaproteobacteria dominate active 2-methyl-4-chlorophenoxyacetic acid herbicide degraders in agricultural soil and drilosphere. Environ Microbiol 13:991–1009.

45. Rodríguez-Cruz MS, Bælum J, Shaw LJ, Sørensen SR, Shi S, Aspray T, Jacobsen CS, Bending GD. 2010. Biodegradation of the herbicide mecoprop-p with soil depth and its relationship with class III *tfdA* genes. Soil Biol Biochem 42:32–39.

46. Becraft ED, Woyke T, Jarett J, Ivanova N, Godoy-Vitorino F, Poulton N, Brown JM, Brown J, Lau MCY, Onstott T, Eisen JA, Moser D, Stepanauskas R. 2017. Rokubacteria: Genomic giants among the uncultured bacterial phyla. Front Microbiol 8:2264.

47. Anantharaman K, Hausmann B, Jungbluth SP, Kantor RS, Lavy A, Warren LA, Rappé MS, Pester M, Loy A, Thomas BC, Banfield JF. 2018. Expanded diversity of microbial groups that shape the dissimilatory sulfur cycle. ISME 12:1715–1728.

48. Hug LA, Thomas BC, Sharon I, Brown CT, Sharma R, Hettich RL, Wilkins MJ, Williams KH, Singh A, Banfield JF. 2016. Critical biogeochemical functions in the subsurface are associated with bacteria from new phyla and little studied lineages. Environ Microbiol 18:159–173.

49. Wang Y, Li T, Li C, Song F. 2020. Differences in microbial community and metabolites in litter layer of plantation and original Korean pine forests in north temperate zone. Microorganisms 8:1–23.

50. Butterfield CN, Li Z, Andeer PF, Spaulding S, Thomas BC, Singh A, Hettich RL, Suttle KB, Probst AJ, Tringe SG, Northen T, Pan C, Banfield JF. 2016. Proteogenomic analyses indicate bacterial methylotrophy and archaeal heterotrophy are prevalent below the grass root zone. PeerJ 4:e2687.

51. Rinke C, Schwientek P, Sczyrba A, Ivanova NN, Anderson IJ, Cheng J-F, Darling A, Malfatti S, Swan BK, Gies EA, Dodsworth JA, Hedlund BP, Tsiamis G, Sievert SM, Liu W-T, Eisen JA, Hallam SJ, Kyrpides NC, Stepanauskas R, Rubin EM, Hugenholtz P, Woyke T. 2013. Insights into the phylogeny and coding potential of microbial dark matter. Nature 499:431–437.

52. Brown CT, Hug LA, Thomas BC, Sharon I, Castelle CJ, Singh A, Wilkins MJ, Wrighton KC, Williams KH, Banfield JF. 2015. Unusual biology across a group comprising more than 15% of domain Bacteria. Nature 523:208–211.

53. Sheremet A, Jones GM, Jarett J, Bowers RM, Bedard I, Culham C, Eloe-Fadrosh EA, Ivanova N, Malmstrom RR, Grasby SE, Woyke T, Dunfield PF. 2020. Ecological and genomic analyses of candidate phylum WPS-2 bacteria in an unvegetated soil. Environ Microbiol 22:3143–3157.

54. Frindte K, Pape R, Werner K, Löffler J, Knief C. 2019. Temperature and soil moisture control microbial community composition in an arctic-alpine ecosystem along elevational and micro-topographic gradients. ISME 13:2031–2043.

55. Ferrari BC, Bissett A, Snape I, van Dorst J, Palmer AS, Ji M, Siciliano SD, Stark JS, Winsley T, Brown M. 2016. Geological connectivity drives microbial community structure and connectivity in polar, terrestrial ecosystems. Environ Microbiol 18:1834–1849.

56. Frey B, Rime T, Phillips M, Hajdas I, Widmer F, Hartmann M. 2016. Microbial diversity in European alpine permafrost and active layers. FEMS Microbiol Ecol 92:18.

57. Pointing SB, Gillor O, Fulthorpe R, Lazzaro A, Hilfiker D, Zeyer J. 2015. Structures of microbial communities in alpine soils: Seasonal and elevational effects. Front Microbiol 6:1330.

58. Nogales B, Moore ERB, Llobet-Brossa E, Rossello-Mora R, Amann R, Timmis KN. 2001. Combined Use of 16S Ribosomal DNA and 16S rRNA to Study the Bacterial Community of Polychlorinated Biphenyl-Polluted Soil. Appl Environ Microbiol 67:1874–1884.

59. Trexler R, Solomon C, Brislawn CJ, Wright JR, Rosenberger A, McClure EE, Grube AM, Peterson MP, Keddache M, Mason OU, Hazen TC, Grant CJ, Lamendella R, Saikaly P, Abdullah K. 2014. Assessing impacts of unconventional natural gas extraction on microbial communities in headwater stream ecosystems in Northwestern Pennsylvania. Front Microbiol 5:1–13.

60. Vallaeys T, Persello-Cartieaux F, Rouard N, Lors C, Laguerre G, Soulas G. 2006. PCR-RFLP analysis of 16S rRNA, *tfdA* and *tfdB* genes reveals a diversity of 2,4-D degraders in soil aggregates. FEMS Microbiol Ecol 24:269–278.

61. Han L, Liu Y, He A, Zhao D. 2014. 16S rRNA gene phylogeny and *tfdA* gene analysis of 2,4-D-degrading bacteria isolated in China. World J Microbiol Biotechnol 30:2567–2576.

62. Park YJ, Kim KH, Han DM, Lee DH, Jeon CO. 2019. *Sphingobium terrigena* sp. Nov., isolated from gasoline-contaminated soil. Int J Syst Evol Microbiol 69:2459–2464.

63. Leys NMEJ, Ryngaert A, Bastiaens L, Verstraete W, Top EM, Springael D. 2004. Occurrence and phylogenetic diversity of *Sphingomonas* strains in soils contaminated with polycyclic aromatic hydrocarbons. Appl Environ Microbiol 70:1944–1955.

64. Niharika N, Moskalikova H, Kaur J, Khan F, Sedlackova M, Hampl A, Damborsky J, Prokop Z, Lal R. 2013. *Sphingobium czechense* sp. nov., isolated from a hexachlorocyclohexane dump site. Int J Syst Evol Microbiol 63:723–728.

65. Li L, Liu H, Shi Z, Wang G. 2013. *Sphingobium cupriresistens* sp. nov., a copper-resistant bacterium isolated from copper mine soil, and emended description of the genus Sphingobium. Int J Syst Evol Microbiol 63:604–609.

66. Chen X, Wang H, Xu J, Song D, Sun G, Xu M. 2016. *Sphingobium hydrophobicum* sp. nov., a hydrophobic bacterium isolated from electronic-waste-contaminated sediment. Int J Syst Evol Microbiol 66:3912–3916.

67. Johnson RC, Deming C, Conlan S, Zellmer CJ, Michelin A, Lee-Lin S, Thomas PJ, Park M, Weingarten RA, Less J, Dekker JP, Frank KM, Musser KA, McQuiston JR, Henderson DK, Lau AF, Palmore TN, Segre JA. 2018. Investigation of a cluster of *Sphingomonas koreensis* infections. N Engl J Med 379:2529–2539.

68. Kantor RS, Miller SE, Nelson KL. 2019. The water microbiome through a pilot scale advanced treatment facility for direct potable reuse. Front Microbiol 10:993.

69. Pinto AJ, Marcus DN, Ijaz UZ, Bautista-de lose Santos QM, Dick GJ, Raskin L. 2016. Metagenomic evidence for the presence of *Comammox Nitrospira*-like bacteria in a drinking water system. mSphere 1:e00054–15.

70. Gomez-Alvarez V, Pfaller S, Revetta RP. 2016. Draft genome sequence of two *Sphingopyxis* sp. strains, dominant members of the bacterial community associated with a drinking water distribution system simulator. Genome Announc 4:e0018316.

71. Crombie AT, Larke-Mejia NL, Emery H, Dawson R, Pratscher J, Murphy GP, McGenity TJ, Murrell JC. 2018. Poplar phyllosphere harbors disparate isoprene-degrading bacteria. PNAS 115:13081–13086.

72. Wu YH, Cheng H, Zhou P, Huo YY, Wang CS, Xu XW. 2015. Complete genome sequence of the heavy metal resistant bacterium Altererythrobacter atlanticus 26DY36T, isolated from deep-sea sediment of the North Atlantic Mid-ocean ridge. Mar Genomics 24:289–292.

73. Hogan DA, Buckley DH, Nakatsu CH, Schmidt TM, Hausinger RP. 1997. Distribution of the *tfda* gene in soil bacteria that do not degrade 2,4-dichlorophenoxyacetic acid (2,4-D). Microb Ecol 34:90–96.

74. Dunning Hotopp JC, Hausinger RP. 2001. Alternative substrates of 2,4-dichlorophenoxyacetate/α-ketoglutarate dioxygenase. J Mol Catal B Enzym 15:155–162.

75. Dunning Hotopp JC, Hausinger RP. 2002. Probing the 2,4-dichlorophenoxyacetate/α-ketoglutarate dioxygenase substrate-binding site by site-directed mutagenesis and mechanism-based inactivation. Biochemistry 41:9787–9794.

76. Amy PS, Schulke JW, Frazier LM, Seidler RJ. 1985. Characterization of aquatic bacteria and cloning of genes specifying partial degradation of 2,4-dichlorophenoxyacetic acid. Appl Environ Microbiol 49:1237–1245.

77. Gonzalez Vazquez AE. 2021. Impact of seasonal environmental variations on 2,4-D fate and metabolism in urban landscapes. PhD Dissertation. University of Wisconsin-Madison, Madison.

78. Davis MW, Jorgensen EM. 2022. ApE, A Plasmid Editor: A freely available DNA manipulation and visualization program. Front Bioinform 2:818619.

79. Tran PQ, Bachand SC, McIntyre PB, Kraemer BM, Vadeboncoeur Y, Kimirei IA, Tamatamah R, McMahon KD, Anantharaman K. 2021. Depth-discrete metagenomics reveals the roles of microbes in biogeochemical cycling in the tropical freshwater Lake Tanganyika. ISME 15:1971–1986.

80. Linz AM, He S, Stevens SLR, Anantharaman K, Rohwer RR, Malmstrom RR, Bertilsson S, McMahon KD. 2018. Freshwater carbon and nutrient cycles revealed through reconstructed population genomes. PeerJ 2018:1–24.

81. Eddy SR. 2011. Accelerated profile HMM searches. PLoS Comput Biol 7:e1002195.

82. Hyatt D, Chen G-L, Locascio PF, Land ML, Larimer FW, Hauser LJ. 2010. Prodigal: prokaryotic gene recognition and translation initiation site identification 111:119.

83. Larsson A. 2014. AliView: A fast and lightweight alignment viewer and editor for large datasets. Bioinformatics 30:3276–3278.

84. Edgar RC. 2004. MUSCLE: A multiple sequence alignment method with reduced time and space complexity. BMC Bioinformatics 5:113.

85. Han L, Liu Y, Li C, Zhao D. 2015. Cloning, expression, characterization and mutational analysis of the *tfdA* gene from *Cupriavidus campinensis* BJ71. World J Microbiol Biotechnol 31:1021–1030.

86. Hogan DA, Smith SR, Saari EA, McCracken J, Hausinger RP. 2000. Site-directed mutagenesis of 2,4-dichlorophenoxyacetic acid/α-ketoglutarate dioxygenase: Identification of residues involved in metallocenter formation and substrate binding. J Biol Chem 275:12400–12409.

87. Stamatakis A. 2014. RAxML version 8: A tool for phylogenetic analysis and post-analysis of large phylogenies. Bioinformatics 30:1312–1313.

88. Gonod LV, Martin-Laurent F, Chenu C. 2006. 2,4-D impact on bacterial communities, and the activity and genetic potential of 2,4-D degrading communities in soil. FEMS Microbiol Ecol 58:529–537.

89. Gazitúa MC, Slater AW, Melo F, González B. 2010. Novel α-ketoglutarate dioxygenase *tfdA-*related genes are found in soil DNA after exposure to phenoxyalkanoic herbicides. Environ Microbiol 12:2411–2425.

90. Cantrell K, Fedarko MW, Rahman G, Mcdonald D, Yang Y, Zaw T, Gonzalez A, Janssen S, Estaki M, Haiminen N, Beck KL, Zhu Q, Sayyari E, Morton JT, Armstrong G, Tripathi A, Gauglitz JM, Marotz C, Matteson NL, Martino C, Sanders JG, Carrieri AP, Song J, Swafford AD, Dorrestein PC, Andersen KG, Parida L, Kim H-C, Vázquez-Baeza Y, Knight R. 2021. EMPress Enables tree-guided, interactive, and exploratory analyses of multi-omic data sets. mSystems 6:1–10.

