## Supplementary Figure 1 and Table 1 for "Expanded diversity of *tfdA* harboring bacteria across the natural and built environment"

[1]Environmental Chemistry and Technology Program
University of Wisconsin – Madison
Madison, WI 53706 USA

[2]Molecular and Environmental Toxicology Program
University of Wisconsin – Madison
Madison, WI 53706 USA

[3]Microbiology Doctoral Training Program
University of Wisconsin – Madison
Madison, WI 53706 USA

[4]Department of Civil and Environmental Engineering
University of Wisconsin – Madison
Madison, WI 53706 USA

[5]Department of Bacteriology
University of Wisconsin – Madison
Madison, WI 53706 USA

* Corresponding author addresses:

Amber M. White: 1550 Linden Drive, Madison WI 53706;; Twitter: @ambermwhite16

Contents (**3 pages**): Figure S1 and Table S1

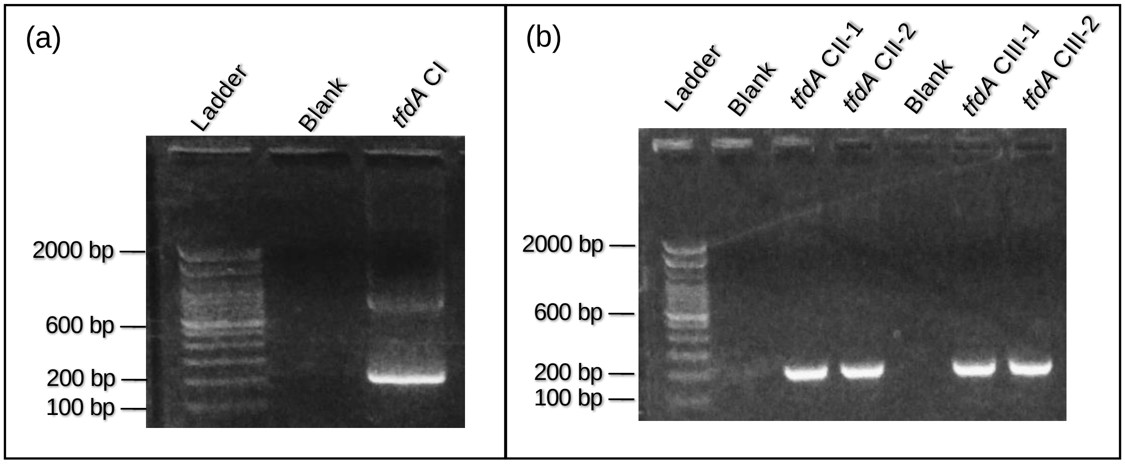


**Figure S1.** Gel visualization of gene product of 215 bp tfdA primer on class I gene standard (a) as well as class II and class III standard (b) in duplicate.

**Table S1.** NCBI BLAST hits for top pairwise alignments between *tfdA* classes I, II, and III genes and sequenced plasmids.

| Reference Gene | Sample | Max Score | Total Score | Query Cover (%) | E-value | Identity (%) |
|---|---|---|---|---|---|---|
| R. eutropha (Class I) | S | 38.3 | 71.1 | 10 | 6.00E-06 | 95.65 |
| | T | 35.6 | 70.2 | 18 | 1.00E-05 | 91.67 |
| | U | 32.8 | 32.8 | 7 | 1.00E-04 | 95.00 |
| | V | 42.8 | 80.1 | 15 | 1.00E-07 | 92.86 |
| | W | 38.3 | 71.1 | 6 | 1.00E-05 | 95.65 |
| B. tropica (Class II) | S | 33.7 | 85.9 | 12 | 7.00E-05 | 91.30 |
| | T | 40.1 | 69.3 | 17 | 1.00E-06 | 95.83 |
| | U | 27.4 | 27.4 | 6 | 6.00E-03 | 94.12 |
| | V | 37.4 | 37.4 | 8 | 4.00E-06 | 92.00 |
| | W | 33.7 | 61.2 | 5 | 1.00E-04 | 91.30 |
| D. acidovorans (Class III) | S | 33.7 | 85.9 | 12 | 7.00E-05 | 91.30 |
| | T | 40.1 | 69.3 | 17 | 1.00E-06 | 95.83 |
| | U | 27.4 | 27.4 | 6 | 6.00E-03 | 94.12 |
| | V | 37.4 | 37.4 | 8 | 4.00E-04 | 92.00 |
| | W | 33.7 | 61.2 | 5 | 1.00E-04 | 91.30 |
